# Phytochrome contributes to blue-light-mediated stem elongation and associated shade-avoidance response in mature *Arabidopsis* plants

**DOI:** 10.1101/2020.11.27.401521

**Authors:** Yun Kong, Youbin Zheng

**Affiliations:** School of Environmental Sciences, University of Guelph, 50 Stone Road East, Guelph, ON N1G 2W1, Canada

**Keywords:** *Arabidopsis*, blue light, flowering time, hypocotyl length, leaf size, phytochrome mutant, shade-avoidance response, stem length

## Abstract

Our recent studies on ornamental plants and microgreens indicate that blue-light-mediated stem elongation is related to phytochrome activity, which was based on the calculated phytochrome photoequilibrium. To examine whether phytochromes really contribute to the blue light’s effect, plant phenotypic responses were investigated in wild type *Arabidopsis* (Col-0), and its quintuple phytochrome (*phyA phyB phyC phyD phyE*) mutant plants under the following light treatments: (1) R, a pure red light from 660-nm LED; (2) B, a pure blue light from 455-nm LED; (3) BR, a impure blue light from LED combination of 94% B and 6% R; and (4) BRF, another impure blue light from LED combination of BR and 6 µmol m^−2^ s^−1^ of FR (735 nm). For all the light treatments, a photosynthetic photon flux density of ≈100 μmol m^−2^ s^−1^ were provided by 24-h lighting daily inside a walk-in growth chamber, which had an air temperature of ≈ 23 °C. The calculated phytochrome photoequilibrium was 0.89, 0.50, 0.69, and 0.60 for R, B, BR, and BRF, respectively, indicating a higher phytochrome activity under R and BR than B and BRF. After 18 days of light treatment, B or BRF increased main stem length in wild-type plants compared with R, but BR had an inhibition effect similar to R. Also, B and BRF relative to R or BR induced earlier flowering and reduced leaf size in wild type plants, showing typical shade-avoidance responses. In phytochrome-deficient mutant plants, the above shade-avoidance responses were inhibited under B or BRF, and induced under BR. However, as an exception, hypocotyl length, a growth trait during the de-etiolation stage, was reduced under B, BR and BRF vs. R regardless of phytochrome absence. It suggests that for mature *Arabidopsis* plants, phytochrome plays an active role in blue-light-mediated stem elongation and associated shade-avoidance response.

## INTRODUCTION

Previous studies using light sources other than light-emitting diode (LED) indicated that blue light (BL), compared with red light (RL), inhibited shoot/leaf elongation (Cosgrove, 1981; Appelgren, 1991; Wheeler et al., 1991; Hoenecke et al., 1992; Brown et al., 1995; Kong et al., 2012). However, in the past decades, studies using LED lighting have reported that stem/leaf elongation was promoted by BL, compared to RL, in some species (Hirai et al., 2006; Hata et al., 2013; Kim et al., 2014; Schwend et al., 2015; Hernandez and Kubota, 2016). Unlike LED, possibly, these non-LED lamps may have provided impure monochromatic light (Bergstrand et al., 2014). For example, the BL from monochromatic fluorescent lamp was reported to contain a low level of other wavelengths, and have a high red/far-red ratio (i.e., 1.87) which may activate phytochromes (Appelgren, 1991).

The promotion effects of pure BL have been also found in our recent studies on ornamental plants and microgreens under LED lighting, and it has been concluded that the promotion effect of pure BL on stem elongation is related to lower phytochrome activity (Kong et al., 2018; Kong et al., 2019a; Kong et al., 2019b;, 2020). In these LED studies, pure BL (B) promoted stem elongation compared with RL (R). However, when a small portion (6 or 10%) of R was added to B, the impure BL (BR) reversed the B promotion effect on elongation, and had similar or greater inhibition effect relative to R. After further adding a low level of far-red light (FR) to BR (R/FR ≈ 1), the resulting impure BL (BRF) recovered the promotion effect similar to B, compared with R. The R/FR reversibility is the classic signature of phytochrome action. Also, as an indicator of phytochrome activity, the phytochrome photostationary state (PPS) value was lower for B (0.49) and BRF (0.63) than R (0.89) and BR (0.74). When the PPS value decreases to 0.60, most plant species show an inactive phytochrome response (Stutte, 2009). Since B reduces PPS below 0.6, possibly the B-promoted elongation is related to low phytochrome activity, and under certain conditions B might need to co-act with R to inhibit elongation growth by increasing phytochrome activity. However, the speculation about the involvement of phytochrome in BL action was only based on reversal response to R/FR and calculated PPS values. It needs further confirmation from direct evidence such as a comparison of phenotypic responses to the above light treatments between wild *Arabidopsis* and its phytochrome mutant plants.

For wild type *Arabidopsis*, co-action with RL was found to increase BL’s inhibition effect on hypocotyl elongation of de-etiolated seedlings in a previous study (Ahmad and Cashmore, 1997). In this study, the inhibitory effect was enhanced by 10 min RL pulses following 10 min BL pulses, but partially reversed by a subsequent 10 min FR pulses. It was concluded that active phytochrome is required for full expression of cryptochrome activity, which mediated BL’s inhibition effect on hypocotyl elongation (Ahmad and Cashmore, 1997). However, differing from our recent study on bedding plants, in this study on *Arabidopsis*, BL alone inhibited stem elongation relative to RL, and co-action with RL only strengthened the BL’s inhibition effect on hypocotyl elongation. The different result about the BL response may be due to different lighting source (non-LED vs. LED), different plant species (*Arabidopsis* vs. bedding plants), and different growth stage (de-etiolation stage vs. vegetative stage). In this case, for mature plants of wild *Arabidopsis* under LED lighting treatments similar to our previous studies, whether B and BRF, relative to R and BR, can promote plant elongation similarly to bedding plants needs further study. Also, phytochrome was only shown to be involved in BL’s inhibition effect on plant elongation in the study by Ahmad and Cashmore (1997). However, it is unknown whether phytochrome also contributes to stem elongation promoted by B or BRF from LED lighting in our previous studies.

In contrast to the above opinion that active phytochrome is required for BL-mediated inhibition effect, some earlier studies on phytochrome-deficient *Arabidopsis* mutants (*phyA* and *phyB*) showed little impairment in BL-dependent inhibition of hypocotyl elongation (Koornneef et al., 1980; Young et al., 1992). However, it has been shown that considerable residual phytochrome responses are retained in all the above phytochrome-deficient mutants (Chory et al., 1989; Reed et al., 1994; Ahmad and Cashmore, 1997). In this case, the possibility that other phytochrome family members (e.g., phyC, phyD and/or phyE) may also contribute to BL-mediated inhibition of hypocotyl elongation cannot be ruled out (Strasser et al., 2010). A recent study on the quintuple phytochrome mutant (*phyA phyB phyC phyD phyE*) indicated that BL alone inhibited hypocotyl elongation of de-etiolated *Arabidopsis* seedlings, suggesting that cryptochrome can operate in the absence of phytochrome (Strasser, et al., 2010). However, in the above studies, the investigation of elongation growth was focused only on hypocotyl length of de-etiolated seedlings and was performed under non-LED lighting which might provide impure BL in many cases. Therefore, the stem elongation response of mature *Arabidopsis* plants needs to be further tested in quintuple phytochrome mutant under BL from LED lighting.

Our recent studies on bedding plants and microgreens indicate that the plant elongation promoted by B or BRF is a shade-avoidance response (Kong, et al., 2018; Kong, et al., 2019b). Besides increased stem elongation, B or BRF, relative to R or BR, caused earlier flowering, smaller cotyledon, longer petiole, and lighter leaf greenness, which varied sensitivity with different species. Possibly, under the same BL treatments (i.e., B or BRF) as our recent study, there is a similar shade-avoidance response in the wild type *Arabidopsis* plants. Since the shade-avoidance response was mediated by BL associated with low phytochrome activity (i.e., B or BRF), it is possible that the quintuple phytochrome mutant may differ from wild type in the response to these BL treatments.

Based on the above information, despite a lot of our previous studies on BL action on stem elongation using LED lighting, the involvement of phytochrome lacked direct proof from the studies on quintuple-phytochrome-deficient mutant plants. In other’s studies on phytochrome-deficient mutants of *Arabidopsis*, the elongation responses to BL have been studied using non-LED lighting and at the de-etiolation stage. For mature *Arabidopsis* plants, the information has so far been unavailable on their stem response to BL (with different PPS values) from LED for both wild type and quintuple phytochrome mutant. The objective of the present study was to examine whether phytochrome contributes to BL-mediated stem elongation as a shade-avoidance response in mature plants by comparing the phenotypic responses to varying-PPS BL between wild *Arabidopsis* and its quintuple phytochrome-deficient mutant (*phyA phyB phyC phyD phyE*) plants under LED lighting.

## MATERIALS AND METHODS

The experiment was performed at the University of Guelph, Guelph, ON, Canada. Two genotypes of *Arabidopsis*, wild type (Col-0) and quintuple phytochrome (*phyA phyB phyC phyD phyE*) mutant (Strasser, et al., 2010), were used for this experiment. Taking into account the low seed germination capacity of this phytochrome-deficient mutant, before seeding, seeds were suspended in GA_4+7_ (Duchefa Biochemie, Haarlem, the Netherlands) solution of 100 μM, and were stratified at 4 °C for 3 d. After rinsing in deionized water three times, the stratified seeds were sown in planting holes (one seed per hole) of a hydroponic system (Fig. 1), with 0.7% Plant Agar (Fisher Scientific, Geel, Belgium), and rockwool cubes (Starter Plugs, Grodan Inc., Ontario, Canada). The two genotypes were evenly and randomly distributed in different rows (i.e., five rows for each genotype) within each tray. The sown trays were placed under the light treatments in a walk-in growth chamber. The ferti-gation method and the environment condition for growing the plants followed the way by Kong and Zheng (2020).

**Fig. 1.**
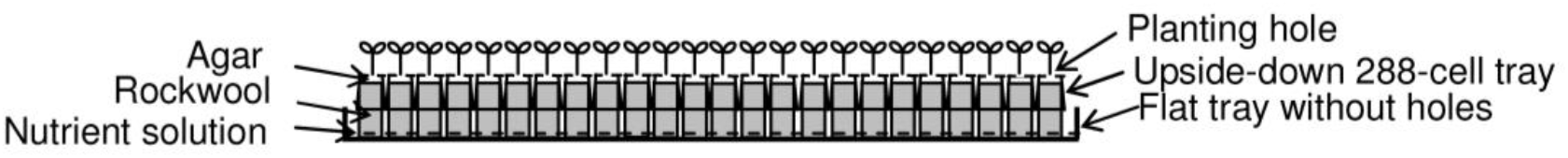
Side-view diagram of a hydroponic system used for growing *Arabidopsis* plants in this experiment.

Light treatments included: (1) R, a pure RL from 660 nm LED; (2) B, a pure BL from 455 nm LED; (3) BR, a impure BL from LED combination of 94% B and 6% R; and (4) BRF, another impure BL from LED combination of BR and 6 µmol m^−2^ s^−1^ of FR (735 nm). Based on the light spectral distribution, the PPS was calculated for each of the four light treatments according to Sager et al. (1988). The calculated PPS values were 0.89, 0.69, 0.60, and 0.50 for R, BR, BRF, and B, respectively. The light treatments were achieved by adjusting the intensities and spectra of a LX602C LED lighting system (Heliospectra AB, Gothenburg, Sweden) using System Assistant 2.0.1 (Heliospectra AB). In the chamber, the four light treatments were arranged to four divided compartments randomly. Opaque curtains were used to separate these compartments to avoid neighboring light pollution. For each light treatment, a photosynthetic photon flux density (PPFD) of around 100 µmol m^−2^ s^−1^ was achieved at the plant canopy level. Light spectra quality and intensity levels were set up and verified for the light treatments using a USB2000+ UV/VIS spectrometer (Ocean Optics, Inc., Dunedin, FL, USA).

Once seed germination was over 50% for each genotype under each light treatment, the cumulative germination percentages were determined. After 18-d lighting, five plants from each genotype in each of light treatments (i.e., one plant from each row in each tray) were randomly selected for investigating plant morphology. The observed plant traits included main stem length, hypocotyl length, rosette leaf number, total leaf number, flowering index, and leaf morphology (size and color). The values of flowering index (ranging from 0–3) were defined as the same as our previous study (Kong and Zheng, 2020). Leaf size and color were observed following the method by Kong et al. (2019b) and Karcher and Richardson (2003).

DPS 7.05 Software (Refine Information Tech. Co., Hangzhou, China), a data processing System, was used for the data analysis. In this experiment, the chamber environment conditions were uniform except for light treatments, and five rows of plants in growing trays were randomly distributed to each combination of light treatments × *Arabidopsis* genotypes. In this case, the experimental arrangement can be considered as a Completely Random Design with two factors and five replicates. Two-way ANOVA was used to determine the effects of each factor (i.e., light treatment, or *Arabidopsis* genotype), and their interaction. For each plant trait, means separation for different treatments were determined using Duncan’s new multiple range test (*P* ≤ 0.05).

## RESULTS

Cumulative germination percentage was not different among the different treatments (data not shown). Under the light treatments, main stem length differed in response pattern between phytochrome mutant and wild type plants (Fig. 2A and B). For wild type plants, B and BRF promoted main stem elongation relative to R or BR, and BR showed an inhibitory effect similar to R, but BRF vs. B had a greater promotion effect. For the phytochrome mutant, plants under B and R showed a similar height, but were shorter than those under BR and BRF, and plants were taller under BR than BRF. Phytochrome mutant had reduced main stem length under B or BRF, but increased main stem length under BR compared to wild type. It suggested that the absence of phytochromes attenuated the enhancement effect of B or BRF and removed the inhibition effect of BR on main stem elongation.

**Fig. 2.**
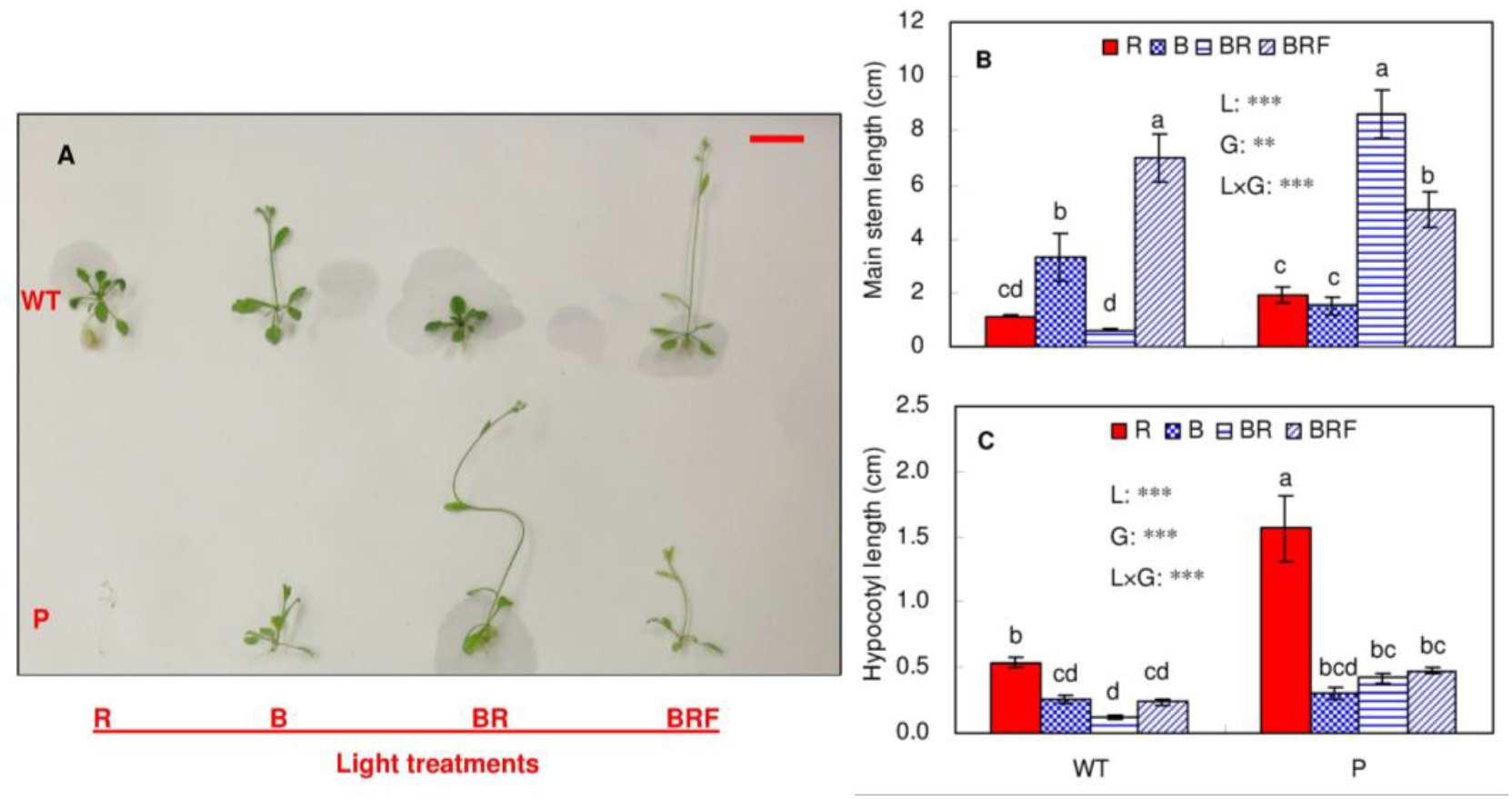
Stem elongation of wild-type *Arabidopsis* and its phytochrome-deficient mutant growing under different light spectra. The picture was taken after 18-d light treatment. The reference bar in the picture is 1.6 cm long. WT = wild type; P = quintuple phytochrome (*phyA phyB phyC phyD phyE*) mutant. For the four light treatments, R = a pure red light from 660 nm LED; B = a pure blue light from 455 nm LED; BR = an impure blue light from LED combination of 94% B and 6% R; and BRF = another impure blue light from LED combination of BR and 6 µmol m^−2^ s^−1^ of FR (735 nm). Data are presented as means ± SE (n = 5). The symbols inside the chart, i.e., L, G and L × G denote light treatment, plant genotype, and their interaction, respectively. Behind the symbols, ns, *, **, or *** indicate no significance or significance at a level of 0.05, 0.01, or 0.001, respectively, for the effect of treatment on the plant trait. Different letters on the data indicate significant difference (Duncan’s new multiple range test, *P* ≤ 0.05).

For hypocotyl length, the light response pattern of phytochrome mutant was similar that of wild type; B, BR, and BRF reduced this trait compared to R, while the three BL treatments were not different from each other (Fig. 2C). The different response in hypocotyl from main stem suggests that BL-mediated elongation growth differed during the early and late growth stages. Under R, phytochrome mutant showed greater hypocotyl length than wild type plants. Hypocotyl was longer under BR for phytochrome mutant than wild type. It suggested that in this case during early growth stage, BL was more effective to inhibit elongation growth than RL, showing an inhibition effect independent of phytochrome.

For total leaf number, the light response pattern of phytochrome mutant was different from that of wild type. B, BR, and BRF, compared to R, did not change total leaf number for wild type, but increased this trait for phytochrome mutant (Fig. 3A). Under R, despite promoting hypocotyl elongation during early growth stage, the quintuple phytochrome mutant was not able to develop beyond some rudimentary leaves at the late stage. This might contribute to the different response of total leaf number between wild and mutant plants.

**Fig. 3.**
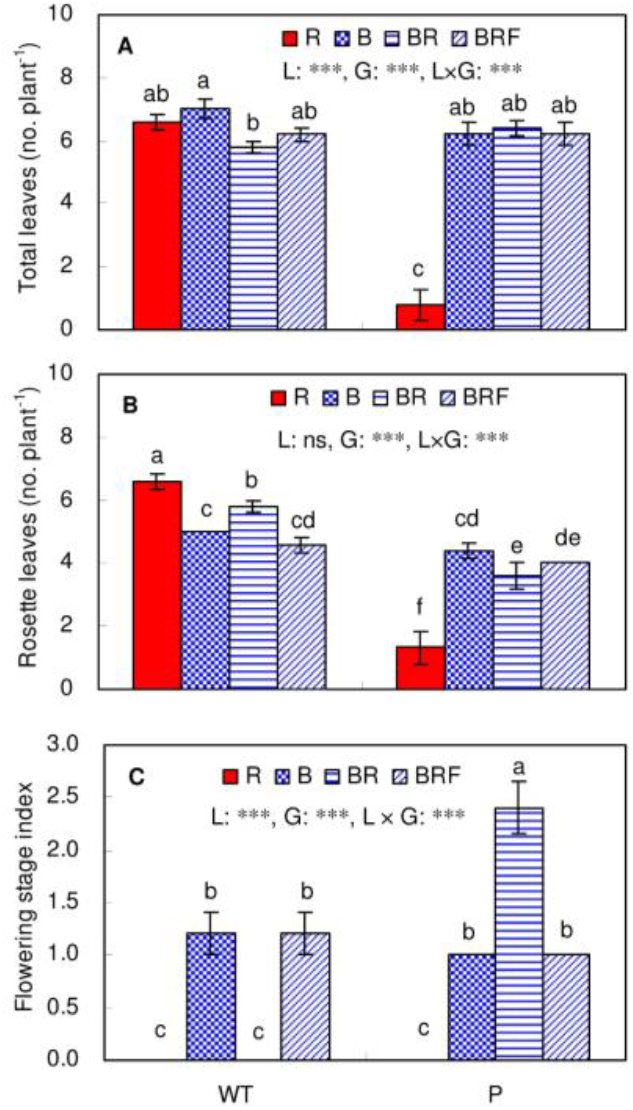
Leaf number and plant flowering of wild-type *Arabidopsis* and its phytochrome-deficient mutant growing under different light spectra. WT = wild type; P = quintuple phytochrome (*phyA phyB phyC phyD phyE*) mutant. For the four light treatments, R = a pure red light from 660 nm LED; B = a pure blue light from 455 nm LED; BR = an impure blue light from LED combination of 94% B and 6% R; and BRF = another impure blue light from LED combination of BR and 6 µmol m^−2^ s^−1^ of FR (735 nm). Data are presented as means ± SE (n = 5). The symbols inside the chart, i.e., L, G and L × G denote light treatment, plant genotype, and their interaction, respectively. Behind the symbols, ns, *, **, or *** indicate no significance or significance at a level of 0.05, 0.01, or 0.001, respectively, for the effect of treatment on the plant trait. Different letters on the data indicate significant difference (Duncan’s new multiple range test, *P* ≤ 0.05).

For rosette leaf number, its light response pattern was similar to total leaf number for the phytochrome mutant, but different from total leaf number for wild type (Fig. 3B). All the BLs (i.e., B, BR, and BRF), relative to R, increased rosette leaf number in phytochrome mutant, but reduced this trait in wild type. For BL, rosette leaves under BR were increased compared with B and BRF for wild-type plants, but were reduced compared with B for phytochrome-deficient mutants. Also, under BR, the phytochrome mutant had less rosette leaves than wild type.

For flowering index, its light response pattern was generally similar to the response of main stem length, but different between wild type and phytochrome mutant (Fig. 3C). In wild type, flowering index was increased under B and BRF, compared with R or BR, but was not different between BR and R, or between BRF and B. In phytochrome mutant, flowering index was increased under B, BR, and BRF compared with R, and the promotion effect was greater for BR than B and BRF, and was similar for B and BRF. Under BR, phytochrome mutant showed a much greater flowering index than wild type. It suggested that for wild *Arabidopsis*, low-PPS BL (i.e., B or BRF) promoted flowering, but high-PPS BL (i.e., BR) inhibited flowering. However, in the absence of phytochromes, BR lost flowering inhibition effect, and promoted flowering to a greater degree than B or BRF.

For petiole length, the light response pattern of wild type was different from that of phytochrome mutant (Fig. 4A). In wild type, BR and BRF reduced petiole length compared to R, but for the phytochrome mutant, BR vs. R increased this trait, and there was no difference between BRF and R. Compared to wild type, phytochrome mutant had a longer petiole under BR, but a shorter petiole under R. This indicated that in the absence of phytochromes that BR vs. R lost the inhibition effect and showed a promotion effect on petiole elongation.

**Fig. 4.**
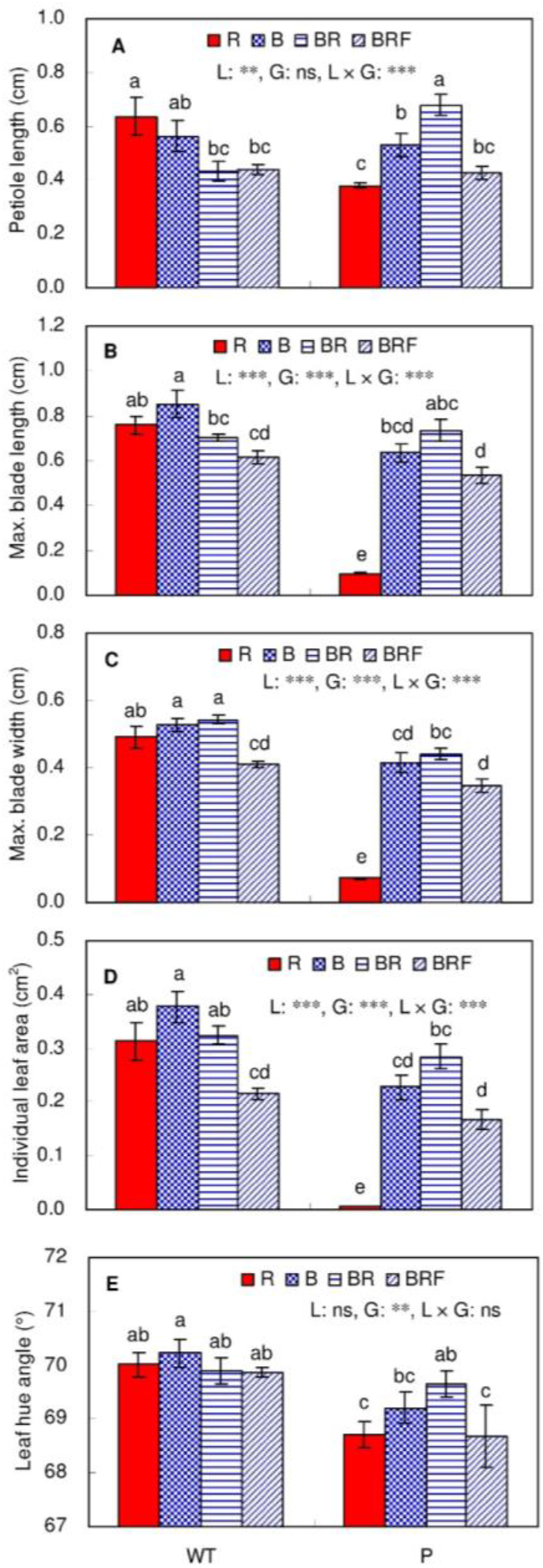
Leaf size and color of wild-type *Arabidopsis* and its phytochrome-deficient mutant growing under different light spectra. WT = wild type; P = quintuple phytochrome (*phyA phyB phyC phyD phyE*) mutant. For the four light treatments, R = a pure red light from 660 nm LED; B = a pure blue light from 455 nm LED; BR = an impure blue light from LED combination of 94% B and 6% R; and BRF = another impure blue light from LED combination of BR and 6 µmol m^−2^ s^−1^ of FR (735 nm). Data are presented as means ± SE (n = 5). The symbols inside the chart, i.e., L, G and L × G denote light treatment, plant genotype, and their interaction, respectively. Behind the symbols, ns, *, **, or *** indicate no significance or significance at a level of 0.05, 0.01, or 0.001, respectively, for the effect of treatment on the plant trait. Different letters on the data indicate significant difference (Duncan’s new multiple range test, *P* ≤ 0.05).

For blade size, and leaf area, the light response pattern of wild type was different from that of phytochrome mutant (Fig. 4B–D). In wild type, BRF, compared to R or B, reduced blade size, and leaf area, but B or BR had similar effects as R on these traits. In the phytochrome mutant, the three traits were increased under B, BR, or BRF relative to R, and were reduced under BRF vs. BR. Under both B and R, phytochrome mutant, compared with wild type, had reduced blade size and leaf area. This indicates that the absence of phytochrome inhibited the leaf expansion under B or R.

For leaf color, the light response pattern of wild type was different from that of phytochrome mutant (Fig. 4E). In wild type, leaf hue angle was not different among the light treatments. In phytochrome mutant, leaf hue angle was increased under BR, compared with R or BRF, but there was no difference among R, B and BRF. Under R, B or BRF, phytochrome mutant, compared wild type, had decreased leaf hue angle. This indicates that in the absence of phytochromes, leaves under R, B or BRF reduced greenness.

## DISCUSSION

### BL’s effect on stem elongation of mature *Arabidopsis* plants is related to phytochrome activity

In wild *Arabidopsis* plants, B and BRF increased main stem length compared with R and BR; the inhibitory effect of BR was similar to that of R, and the promotion effect of BRF was greater than that of B. Also, as an indicator of phytochrome activity, the PPS value is lower for B (0.50) and BRF (0.60) than BR (0.69) and R (0.89). This suggests that BL’s effect on stem elongation is related to phytochrome activity, which is similar to the elongation response to BL found in our previous study on bedding plants (Kong, et al., 2018). However, unlike bedding plants, in wild type *Arabidopsis*, BRF promoted stem elongation to a larger degree compared to B, rather than a similar promotion effect (Kong, et al., 2018). It appeared that BRF had an additive promotion effect of B and FR on stem elongation in the wild type *Arabidopsis* plants. The underlying mechanism needs further study for the differences in response to B and BRF among plant species.

Unlike main stem, the above elongation response to BL did not occur in the hypocotyl. For hypocotyl length, all the BL treatments (B, BR, and BRF) showed similar inhibitory effects compared to R. It appeared that in the context of current study, BL-mediated hypocotyl elongation was affected little by phytochrome activity. Similar inhibitory effects of BL vs. RL on hypocotyl elongation has been found in a previous study on *Arabidopsis* (Ahmad and Cashmore, 1997). However, differing from our current study, the BL’s inhibition effect was strengthened when followed by a RL pulse and weakened when followed by a FR pulse, suggesting the contribution of phytochrome activity to BL-mediated hypocotyl elongation. The inconsistency of the results may be partly explained by the different light intensity employed in the two studies: a much lower BL intensity was used in the previous study (≈ 30 µmol m^−2^ s^−1^) than in our present study (≈ 100 µmol m^−2^ s^−1^). At least for some species, inhibitory effect on elongation by either pure or impure BL strengths with light intensity increasing (Cope and Bugbee, 2013; Johnson et al., 2019). In this case, regardless of phytochrome activity, BL at a PPFD of 100 µmol m^−2^ s^−1^ might be strong enough to inhibit hypocotyl elongation in de-etiolated seedlings, rather than stem elongation in mature plants for *Arabidopsis*.

Although the PPS value, phytochrome photoequilibrium calculated according to absorption of light spectrum, can be easily used to indicate indirectly the phytochrome activity under different BL treatments, it may not reflect the real situation. The absorption of light spectrum was measured in the solution with isolated and purified phytochrome, rather than in plant leaves (Sager, et al., 1988). Masking pigments (predominantly chlorophyll), and leaf structure can alter intracellular light regimes around phytochrome (Gardner and Graceffo, 1982). Studying the difference between phytochrome mutants and wild type is another way to explore the involvement of phytochrome in BL-mediated elongation growth. However, many early studies on *phyA phyB* mutants cannot exclude the involvement of other residual phytochrome species (Strasser, et al., 2010). In this case, quintuple phytochrome mutant, which is deficient of all the currently known phytochrome species, may provide a new plant material to study the mechanism of BL’s action on stem elongation.

### Phytochrome play an active role in BL-mediated stem elongation of mature plants in *Arabidopsis*

The pattern of main stem length response to the BL treatments was totally different between the phytochrome mutant and the wild *Arabidopsis* in the present study. For plants under the light treatments following an order of, R, B, BR, and BRF, main stem was short-tall-short-tall for wild type, but was short-short-tall-short for phytochrome mutant. The different response between the wild type and phytochrome mutant suggests that phytochrome is actively involved in the BL-mediated main stem elongation. Recent studies on *Arabidopsis* indicates that transcriptional changes in response to BL can be coordinately regulated by a cross talk at least between cryptochrome and phytochrome due to some of the shared signaling pathways (Liu et al., 2016; Pedmale et al., 2016; Mishra and Khurana, 2017; Su et al., 2017; Yang et al., 2017). Possibly, phytochrome activity can modify the function of cryptochrome, the BL receptor, on main stem elongation (Liu, et al., 2016).

Obviously, the above difference in main stem length between wild type and phytochrome mutant plants resulted from their different responses to each of the three BL treatments. Under BR (i.e., BL with a higher PPS value), main stem was the tallest for phytochrome mutant, but was the shortest for wild type among the light treatments. The reversal effect of BR on elongation in the presence or absence of phytochrome indicated that active phytochrome played an important role in the inhibitory effect of BR on stem elongation for wild *Arabidopsis*. Under B or BRF (i.e., BL with lower PPS values), phytochrome mutant reduced main stem length compared to wild type. In the absence of phytochrome, the promotion effect on main stem elongation was eliminated under B and reduced under BRF relative to R. It appeared that weak-activity phytochrome might contribute partly to increased main stem elongation under BL associated with low PPS for wild types. Possibly, some other photoreceptors (e.g., phototropins), in addition to phytochromes, were also partly involved in the BL-promoted elongation (Kong and Zheng, 2020).

Differing from main stem length, hypocotyl length was reduced under B, BR, and BRF relative to R for both phytochrome mutant and wild type, showing a similar response pattern between the two *Arabidopsis* genotypes. It appears that phytochrome is not required for cryptochrome to inhibit hypocotyl elongation under BL in some cases (Strasser, et al., 2010). It is well known that hypocotyl elongation occurs only at early growth stage, but main stem elongation lasts until late growth stage. Possibly, the involvement of phytochromes in the BL-mediated elongation was less active during early vs. late growth stage for *Arabidopsis* under the conditions (e.g., ≈ 100 µmol m^−2^ s^−1^) in the present study. However, the BL’s inhibition effect on hypocotyl length was greater for phytochrome mutant than wild type. This was mainly due to the failed inhibition of hypocotyl elongation growth by R for the phytochrome mutant rather than wild type. Similar stretching hypocotyl response to RL has been found in *phyA phyB* double mutant of *Arabidopsis* (Reed, et al., 1994), *phyA phyB phyC* triple mutant of rice (Takano et al., 2009), and quintuple phytochrome mutant of *Arabidopsis* (Strasser, et al., 2010).

### BL-promoted elongation growth of mature *Arabidopsis* plants is a shade-avoidance response partly mediated by phytochrome

In *Arabidopsis*, shade signals cause great changes in plant growth and morphology, which are called shade-avoidance responses, including promoted elongation of stems and petioles, reduced leaf size, and enhanced early flowering (Casal, 2012). In the current study, for the wild type plants, B or BRF, compared with R or BR, not only increased main stem length, but also reduced rosette leaf number and increased flowering index indicating an earlier flowering. Moreover, leaf size (including leaf area, and maximum blade length and width) was reduced under BRF vs. R, despite varying to a much smaller degree than flowering index and main stem length. This suggests that the plant elongation promoted by B or BRF is a shade-avoidance response in wild type *Arabidopsis*. Similar shade-avoidance responses have been also observed under B or BRF in our recent study on bedding plants (Kong, et al., 2018).

It is worthwhile to note that, for wild type *Arabidopsis*, plants under B or BRF relative to R or BR did not show shade-avoidance responses in some traits such as hypocotyl length, petiole length, and leaf color. It appeared that for the same plant, different plant traits had varied sensitivity in shade-avoidance response to BL with low PPS. This was supported by our previous studies on other plant species (Kong, et al., 2018; Kong, et al., 2019b). Hypocotyl length and leaf morphology showed a lower sensitivity than main stem length and flowering index. This possibly resulted from varied threshold levels for BL to induce the shade-avoidance response at different stages or in different cells (Mishra and Khurana, 2017). In other words, threshold level of BL to prevent shade-avoidance response may be lower at early vs. late growth stage, and in leaves vs. stems. Also, it implies that only under suboptimal, rather than optimal, light levels, can BL effects on these traits be affected by phytochrome activity, i.e., there is a co-action between cryptochromes and phytochromes (Casal and Mazzella, 1998)

Differing from wild type plants, phytochrome-deficient plants showed some antagonized shade-avoidance responses under B or BRF rather than BR or R. For phytochrome mutant, B or BRF increased leaf size relative to R, and increased rosette leaf number, and reduced flowering index, petiole length, and main stem length relative to BR. Apparently, for phytochrome mutant, shade-avoidance responses were induced under R or BR, but were prevented under B or BRF. This was contrasting to the shade-avoidance responses observed under B or BRF for wild type plants. It appears that quintuple phytochrome mutant can change the shade-avoidance responses induced in wild types under B or BRF relative to BR or R. However, phytochrome mutant showed shade-avoidance responses under R mainly at early growth stage, and under BR mainly at late growth stage, because phytochrome mutant exposed to R rather than BR had arrested development after the cotyledon stage (Strasser, et al., 2010). Nevertheless, it suggests that phytochromes play an important role in the prevention of shade-avoidance response by R and BR. However, even in absence of phytochromes, some shade-avoidance responses (e.g., reduced leaf size and greenness) were still found under BRF vs. BR. This suggests that some other photoreceptors, in addition to phytochrome, might be partly involved in the shade-avoidance response of wild *Arabidopsis* induced by BL with low PPS.

Overall, for wild type *Arabidopsis* plants, BL with low PPS (i.e., B or BRF), relative to R, promoted main stem elongation, but BL associate with high PPS (i.e., BR) showed a similar inhibition effect as R. The absence of phytochrome reduced and even eliminated the promotion effect of B and BRF, and reversed BR effect from inhibition to promotion. However, B, BR and BRF vs. R reduced hypocotyl length in both wild types and phytochrome mutants. This suggests that in mature *Arabidopsis* plants, BL’s effect on stem elongation is related to phytochrome activity, and phytochrome is actively involved in BL-mediated stem elongation. Along with enhanced main stem elongation, B and BRF, compared with R or BR, also induced earlier flowering and reduced leaf size in wild type plants, suggesting that the B- or BRF-promoted stem elongation is one of shade-avoidance responses. In the absence of phytochrome, the above shade-avoidance responses were inhibited under B or BRF, and induced under BR. Therefore, phytochrome contributes to BL-mediated stem elongation and associated shade-avoidance response in mature *Arabidopsis* plants.

## ACKNOWLEDGEMENTS

This research work was supported by grants from Natural Sciences and Engineering Research Council of Canada. The LED lamps used for this study were provided by Heliospectra AB (Gothenburg, Sweden). We are grateful to Dr. Pablo D. Cerdána and Dr. Maximiliano Sánchez-Lamasa for their donation of quintuple phytochrome mutant seeds, and instruction of seed germination promotion. We also thank Dr. Yongmei Bi for her donation of wild-type *Arabidopsis* and instruction of cultivation technique. Thanks also go to Katherine Schiestel and Dave Llewellyn for their excellent technical supports during the trial and Devdutt Kamath for his help with editing the manuscript.

## AUTHOR CONTRIBUTIONS

YK designed the experiment, and collected, analyzed, and interpreted the data, and drafted and revised the manuscript. YZ participated in the whole process of this study as the principal investigator. Both authors approved the final manuscript.

## CONFLICT OF INTEREST STATEMENT

No conflict of interest.

## Notes

### Competing Interest Statement

The authors have declared no competing interest.

